# Dissociating the incubation of appetitive and consummatory behavior in a model of oral cocaine self-administration

**DOI:** 10.1101/2025.02.12.637922

**Authors:** Mark D. Namba, Samuel L. Goldberg, Christine M. Side, Abhiram Yadlapalli, Christina M. Curran-Alfaro, Devin Kress, Melissa F. Fogarty, Amanda L.A. Mohr, Jacqueline M. Barker

## Abstract

Cocaine use disorder remains a persistent public health dilemma that currently lacks effective treatment strategies. One key impediment to successful treatment outcomes is increased drug craving that occurs over the course of abstinence and subsequent relapse to drug use. This phenomenon, known as the incubation of drug craving, has been modeled extensively in rodent models of intravenous drug self-administration. As commonly implemented, the design of intravenous self-administration preclinical studies precludes disentangling appetitive and consummatory behaviors as drug seeking (appetitive) and taking (consummatory) is simultaneous. Here, we employed a model of oral cocaine self-administration to interrogate the incubation of drug vs nondrug craving, where the route of administration is identical between reinforcers and appetitive and consummatory behaviors are dissociable. Oral self-administration of cocaine produced detectable levels of cocaine and its metabolite, benzoylecgonine, within the blood and brain, and blood and brain levels of both substrates correlated with cocaine consumption. When tested for seeking-(lever pressing) and taking-related (magazine head entries) behavior after 1 or 21 days of forced abstinence from cocaine or saccharin, we observed incubation of lever pressing among cocaine-administering mice and incubation of magazine entries among saccharin-administering mice. These behavioral changes were accompanied by reduced expression of the glial glutamate transporter GLT-1 within the nucleus accumbens (NAc) of cocaine self-administering mice, regardless of abstinence. Altogether, these results underscore the utility of this model of cocaine self-administration, highlight the conserved nature of incubated cocaine seeking across routes of administration, and demonstrate the dissociable neurobehavioral sequelae of the incubation of reward seeking across reinforcer types.

## Introduction

Over the last decade, cocaine use and overdose deaths have risen sharply across the world, especially within the United States (Mattson et al., 2021). Unlike other addictive drugs, there are no FDA-approved medications available for the treatment of cocaine use disorders (CUDs), which directly contributes to this significant public health dilemma. A major barrier to successful CUD treatment outcomes is drug relapse, which is often precipitated by exposure to drug-associated contexts and environmental cues that can trigger intense cravings and activation of corticolimbic substrates that mediate drug-seeking behavior (Carter and Tiffany, 1999; Childress et al., 1999; Parvaz et al., 2016). In both humans and animals, and across drug classes, drug craving and seeking behavior increases over the course of a protracted abstinence period (Pickens et al., 2011; Li et al., 2015). Thus, identifying neural mechanisms that facilitate this time-dependent increase in drug seeking represents a promising avenue for pharmacotherapeutic development.

A major limitation of the extant preclinical literature on the incubation effect is that many of these studies demonstrate this effect following intravenous drug self-administration (Grimm et al., 2003; Conrad et al., 2008; Abdolahi et al., 2010; Pickens et al., 2011; Venniro et al., 2016). An inherent assumption that is borne out of these studies is that the incubation effect observed at the lever that previously delivered drug (i.e., the “active lever”) represents a potentiation of an appetitive cocaine *seeking* process (driven by drug-associated cues). However, in the case of intravenous drug self-administration (particularly on a FR1 schedule), drug *seeking* and *taking* are inherently conflated and not dissociable (Roberts et al., 2013). Thus, it remains unclear whether the incubation effect reported in the preclinical literature represents a time-dependent enhancement of appetitive or consummatory behavior and whether the route of administration impacts this process. Beyond this, many self-administration studies in mice are restricted in their scope by the known practical issues regarding intravenous drug self-administration in mice compared to rats (e.g., poorer catheter patency). This further highlights the utility of alternative self-administration models that may be more amenable to studies that utilize mice (e.g., transgenic mouse models and/or adolescent mice), require longitudinal experimental timelines that are not compatible with long-term intravenous catheter patency, include biohazardous models that make catheter surgery impractical and unsafe (e.g., HIV-infected humanized mice), among many others. Indeed, previous studies have utilized oral self-administration of cocaine in an operant setting to examine the neurobehavioral sequelae of cocaine seeking (Miles et al., 2003; Depoy et al., 2016; Li et al., 2022). Taken altogether, examining the incubation of craving across varying drug access conditions represents an important and necessary step forward in our understanding of the neurobehavioral underpinnings of this critical drug relapse-associated process.

Corticolimbic glutamatergic plasticity is a key neurobiological adaptation that underlies the incubation of drug craving and cue-induced drug seeking (Cornish and Kalivas, 2000; Kalivas and McFarland, 2003; McFarland et al., 2003; Kalivas, 2009; Pickens et al., 2011; Wolf and Tseng, 2012; Ma et al., 2014). Specifically, changes in the synaptic expression of AMPA receptor subunits within the nucleus accumbens (NAc) are associated with enhanced drug-seeking behavior. For example, increased expression of the AMPA receptor (AMPAR) subunit GluA1, which confers AMPAR calcium permeability, is observed within the NAc after chronic cocaine self-administration and protracted abstinence (Conrad et al., 2008; Siemsen et al., 2019). Astrocytes play a crucial role in regulating AMPA receptor-mediated glutamatergic transmission within the NAc and subsequent cocaine-seeking behavior. The glial glutamate transporter GLT-1 (also known as EAAT-2) mediates astrocytic uptake of extracellular glutamate (Kalivas, 2009; Miguel-Hidalgo, 2009; Scofield and Kalivas, 2014; Roberts-Wolfe and Kalivas, 2015). Cocaine self-administration impairs the function and expression of GLT-1 within the NAc, and restoration of these deficits suppresses cue-induced cocaine seeking (Knackstedt et al., 2010; Trantham-Davidson et al., 2012; Reissner et al., 2015).

In this this study, we interrogated the incubation of cocaine seeking in a model of operant, oral cocaine self-administration in male mice and probe the effects of this model of cocaine self-administration on NAc glutamatergic substrates. A key finding of this study is that mice self-administering cocaine exhibit an incubation of seeking behavior (lever pressing) after protracted abstinence, while mice self-administering saccharin exhibit incubation of taking-related (magazine entry) behavior. Concomitantly, we observed a downregulation of GLT-1 expression within the NAc following cocaine self-administration. Together, these findings inform the neurobehavioral mechanisms that underlie the expression of incubated drug craving and relapse, which is vital for identification of effective pharmacotherapeutic strategies for treating substance use disorders (SUDs).

## Methods

### Subjects

Nine week-old male C57BL/6J mice (N=57) from Jackson Laboratories (Bar Harbor, ME, USA) were housed in, and acclimated to, a temperature-and humidity-controlled vivarium with *ad libitum* access to standard chow and water for 5 days prior to behavioral testing. Throughout behavioral testing, mice were food restricted to 90% of their free-feeding body weight. All procedures were approved by the Drexel University Institutional Animal Care and Use Committee and adhered to the NIH’s *Guide for the Care and Use of Laboratory Animals*.

### Drugs & key reagents for behavioral studies

Saccharin sodium dihydrate (Spectrum Chemical MFG, New Brunswick, NJ, USA) was dissolved in tap water to a final concentration of 0.1% w/v. Cocaine hydrochloride (NIDA Drug Supply Program, RTI International, Research Triangle Park, NC, USA) was dissolved in 0.1% saccharin solution to a concentration of 75 or 100 µg/mL. These concentrations were based on a previous study (Depoy et al., 2016) and pilot experiments from our lab demonstrating that increasing the concentration beyond 100 µg/mL does not improve cocaine consumption or seeking behavior (**Supp. Fig S1**).

### Cocaine and saccharin self-administration

Behavioral training took place in operant conditioning chambers containing two levers (one active, one inactive), a syringe pump, a magazine for liquid reward consumption, a house light, and a tone generator (Med Associates). Prior to standard 2-hour self-administration training sessions, mice received an overnight training session in which they self-administered a 0.1% saccharin solution on a FR1 schedule of reinforcement (20 µL/reinforcer), where light and tone cues were paired with the duration of saccharin delivery. This was performed to facilitate the acquisition of self-administration. After a rest day, mice were trained to self-administer saccharin (FR1 schedule, 50 µL/reinforcer) for 3 days for 2 hours per session. Mice in the saccharin control group continued to self-administer saccharin for another 15 days, while mice in the cocaine group were transitioned to self-administration of 75 µg/mL cocaine + 0.1% saccharin for 3 days followed by 100 µg/mL cocaine + 0.1% saccharin for 12 days. Total lever presses, magazine entries, and reinforcers earned were analyzed between cocaine-and saccharin self-administering mice as well as magazine checking behavior, which is defined as the percentage of lever presses that are immediately followed by a magazine entry.

### Toxicology standards and reagents

Cocaine, benzoylecgonine, cocaine-d_3_, and benzoylecgonine-d_3_ were all obtained from Cerilliant Corporation (Round Rock, TX, USA). Formic acid ampoules were obtained from Life Technologies Corporation (Carlsbad, CA, USA). Liquid chromatography mass spectrometry (LCMS) grade water and acetonitrile were obtained from Honeywell (Muskegon, MI, USA). Ammonium formate was obtained for Millipore Sigma (St Louis, MO, USA). Sodium borate decahydrate was obtained from Fisher Scientific (Waltham, MA, USA). Methyl-tert-butyl ether (MTBE), 2-propanol, and defibrinated sheep blood were all obtained from VWR (Radnor, PA, USA).

### Cocaine and metabolite assessments from blood and brain tissue samples

One cohort of mice was trained to self-administer cocaine as described above, and all mice were briefly anesthetized with 2-5% isoflurane for collection of whole submandibular blood at two different timepoints (in a counterbalanced manner) – after 1 or 2 hours of self-administration. After the final blood collection timepoint, mice were deeply anesthetized with isoflurane and sacrificed via rapid decapitation for brain tissue collection. Brains were frozen over dry ice and stored at -80°C until later processed for cocaine and BZE analysis. A timeline of these experiments is provided in **Figure 1A** and self-administration behavior for these animals is depicted in **Supplemental Figure S2**. Samples were analyzed using a Waters Acquity UPLC coupled to a Waters Xevo TQ-S Micro triple quadrupole mass spectrometer (LC-MS-MS). Chromatographic separation was achieved using on an Agilent Poroshell C18 column (3μm, 2.7x100mm) with 5mM ammonium formate (pH 3) (MPA) and 0.1% formic acid in acetonitrile (MPB) with a flow rate of 0.4 mL/minute with a 5 µL injection volume. The gradient for the method can be found in **Supplemental Table 1**.

**Figure 1.**
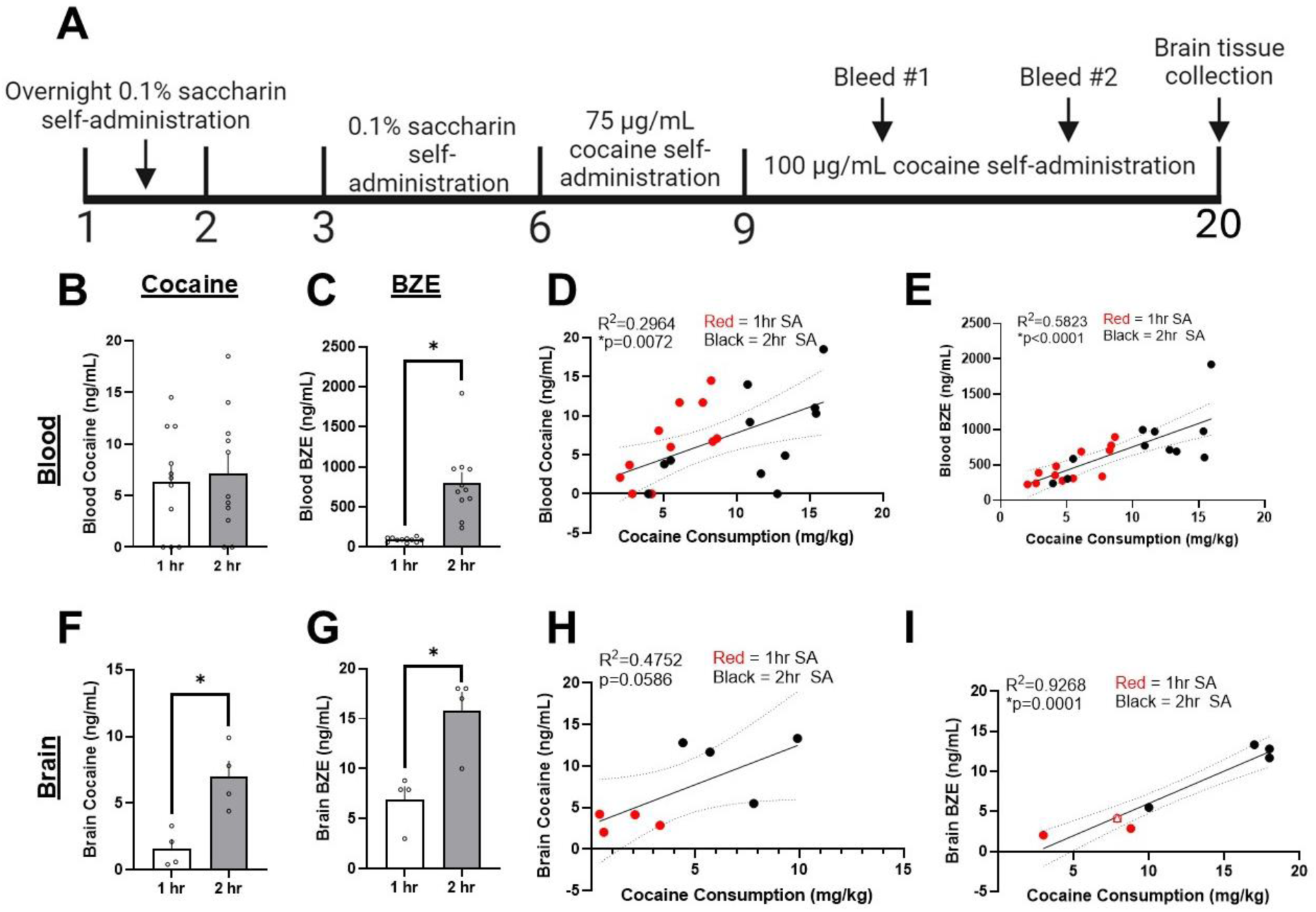
Timeline of cocaine self-administration procedures and tissue concentrations of cocaine and benzoylecgonine (BZE) following cocaine self-administration. (A) Mice were trained in an overnight session of 0.1% saccharin self-administration at 20 µL/reinforcer prior to self-administration of 50 µL/reinforcer of saccharin for 3 days. Mice then self-administered 75 µg/mL cocaine for 3 days followed by 100 µg/mL cocaine for 10 days. Submandibular blood was collected after 1 or 2 hours of self-administration at sessions 3 and 8 of 100 µg/mL cocaine self-administration and were sacrificed for brain tissue collection at session 11 once cocaine consumption recovered. (B) Blood cocaine did not significantly differ between 1 and 2 hours of cocaine self-administration. (C) A significant increase in BZE was observed after 2 hours of self-administration compared to 1 hour. (D-E) A significant correlation was observed between cocaine consumption (mg/kg) and observed levels of blood cocaine and BZE. (F) A significant increase in brain cocaine levels was observed after 2 hours of cocaine self-administration compared to 1 hour. (G) Similarly, a significant increase in brain BZE levels was observed after 2 hours of cocaine self-administration compared to 1 hour. (H-I) While a trend towards a significant correlation between cocaine consumption and brain cocaine levels was observed (p = 0.0586), a significant correlation between cocaine consumption and brain BZE levels was observed. *p < 0.05. Black dots = 1 hour self-administration. Red dots = 2 hour self-administration. *Note: The open circle and open triangle data points in Figure I are stacked because X/Y coordinate values were too close to discriminate visually*.

A liquid-liquid extraction (LLE) was developed for the analysis of cocaine and BZE using 75 µL of blood. After aliquoting, calibrators and controls received 25 uL of the corresponding 50:50 methanol:water standard drug solution, and blanks received 50:50 methanol:water, for all samples to be normalized to 100 μL. 25 µL of internal standard, 100 µL of 0.1M bicarbonate buffer (pH = 9) and 1 mL of 70:30 MTBE:2-propanol were added. Samples were capped and rotated for five minutes followed by centrifugation at 4000 rpm for four minutes. Samples were frozen at -80°C, and the organic layer was transferred to a new tube. Samples were dried to completion at 45°C and reconstituted in 200 µL of 90:10 MPA:MPB.

Quantitative method verification was performed in-house prior to sample analysis. Parameters evaluated included calibration model, bias and precision, limit of quantification (LOQ), interferences, and matrix effects. The calibration curve for cocaine was 2, 5, 25, 125, 250, 625, and 1250 ng/mL with QC’s at 10 and 1000 ng/mL, and for benzoylecgonine it was 50, 100, 250, 500, 1000, 2500, and 5000 ng/mL with QC’s at 200 and 4000 ng/mL. The limit of detection was administratively set at 2 ng/mL for cocaine and 50 ng/mL for benzoylecgonine.

For brain samples, standard addition was used to calculate concentration. This process involved taking four 100 µL aliquots per sample. Three of the four replicate samples were fortified with the analytes of interest, termed “up-spikes” to generate an internal calibration curve. A preliminary analysis was performed on the brain samples following the blood procedure described above to determine the starting concentrations of these samples. The resulting concentration determined the unique “up-spike” values and if dilution was needed. Internal standard was added to all samples prior to extraction via LLE. Following LC–MS-MS analysis, resulting analyte-internal standard peak area ratios were plotted against the up-spike concentrations. A linear fit between all data points was implemented. Correlation between data points was assessed (R^2^ > 0.98). The concentration of each analyte in the sample was determined by calculating the y-intercept of the plotted line.

### Forced abstinence and assessment of incubated seeking behavior

Following self-administration, a separate cohort of mice were placed into 1 or 21 days of forced abstinence prior to a 1-hour cue test, where active lever presses resulted in presentation of reward-paired cues but no reward delivery into the magazine. Total lever presses, magazine entries, and magazine checking behavior (as described above) were assessed. Mice were euthanized via rapid decapitation immediately after cue testing for brain tissue collection.

### Cellular subfractionation and western blotting

NAc tissue lysates from mice that underwent forced abstinence and cue testing were prepared for crude membrane (P2) fractions as described previously (Namba et al., 2020). Briefly, tissue punches were homogenized in an ice-cold 0.32M sucrose buffer solution containing protease and phosphatase inhibitors prior to centrifugation at 1,000 x g for 10 mins at 4C. The resulting pellet (i.e., nuclear fraction) was discarded and the supernatant was centrifuged at 12,000 x g for 20 mins at 4C. The resulting pellet was briefly washed with sucrose buffer and then resuspended in 50 µL of 1X PBS containing 1% SDS. Protein concentrations from these P2 homogenates were determined using the Pierce BCA Assay Kit (Thermo Scientific, Waltham, MA, USA) according to manufacturer instructions prior to western blotting. Western blots were performed as previously described (Namba et al., 2024). Briefly, equal microgram quantities of protein (2 µg/well) were separated on 4-12% bis-tris gels and transferred to PVDF membranes, which were blocked in 5% non-fat dried milk in 1X tris-buffered saline + 0.1% Tween-20 (1x TBST) for 2 hours prior to overnight incubation in primary antibodies (see **Supplemental Table 2**). Blots were washed 3 x 10 mins in blocking buffer and probed with HRP-conjugated secondary antibody (**Supp. Table 2**) for 1 hour and then washed in 1x TBST for 6 x 5 mins. Membranes were activated for 5 mins with SuperSignal™ West Pico PLUS Chemiluminescent Substrate (Thermo Scientific, Waltham, MA, USA) and developed using a LiCor digital imager. Band intensities were determined using ImageJ. For every probed target, band intensities were first normalized to that of a standard sample that was used across all gels, and these values for every protein target of interest were then normalized to that of an internal loading control (GAPDH).

### Data analyses

Blood and brain concentrations of cocaine and BZE were analyzed via two-tailed paired (blood) or unpaired (brain) t-tests, and correlations with cocaine consumption were analyzed via simple linear regression. Cocaine and saccharin self-administration behavior was analyzed via two- and three-way repeated measures ANOVAs, with lever (active or inactive), reinforcer (cocaine or saccharin), and session as factors. Active lever presses and magazine entries during the post-abstinence cue test were analyzed via two-way ANOVAs, with abstinence (1d or 21d ABS) and reinforcer (cocaine or saccharin) as factors. Protein expression from western blots was similarly analyzed via two-way ANOVAs. Magazine checking behavior during self-administration and the cue test, defined as the percentage of active lever presses that were immediately followed by a magazine entry, were analyzed via two-tailed unpaired t-test and two-way ANOVA, respectively. NAc protein expression was analyzed via two-way ANOVAs and correlations between protein expression and active lever pressing or magazine entries during the cue test were analyzed via simple linear regression. Holm-Šídák corrections were used for *post hoc* statistical comparisons. Statistical outliers were analyzed via the ROUT method (Q = 1%), which resulted in two statistically significant outliers being removed from the NAc GLT-1 analysis. Geisser-Greenhouse corrections were applied to analyses where the assumption of sphericity was violated (specifically Figure 2). For all analyses, the threshold for statistical significance was set at α = 0.05. All analyses were performed using GraphPad Prism v10.

**Figure 2.**
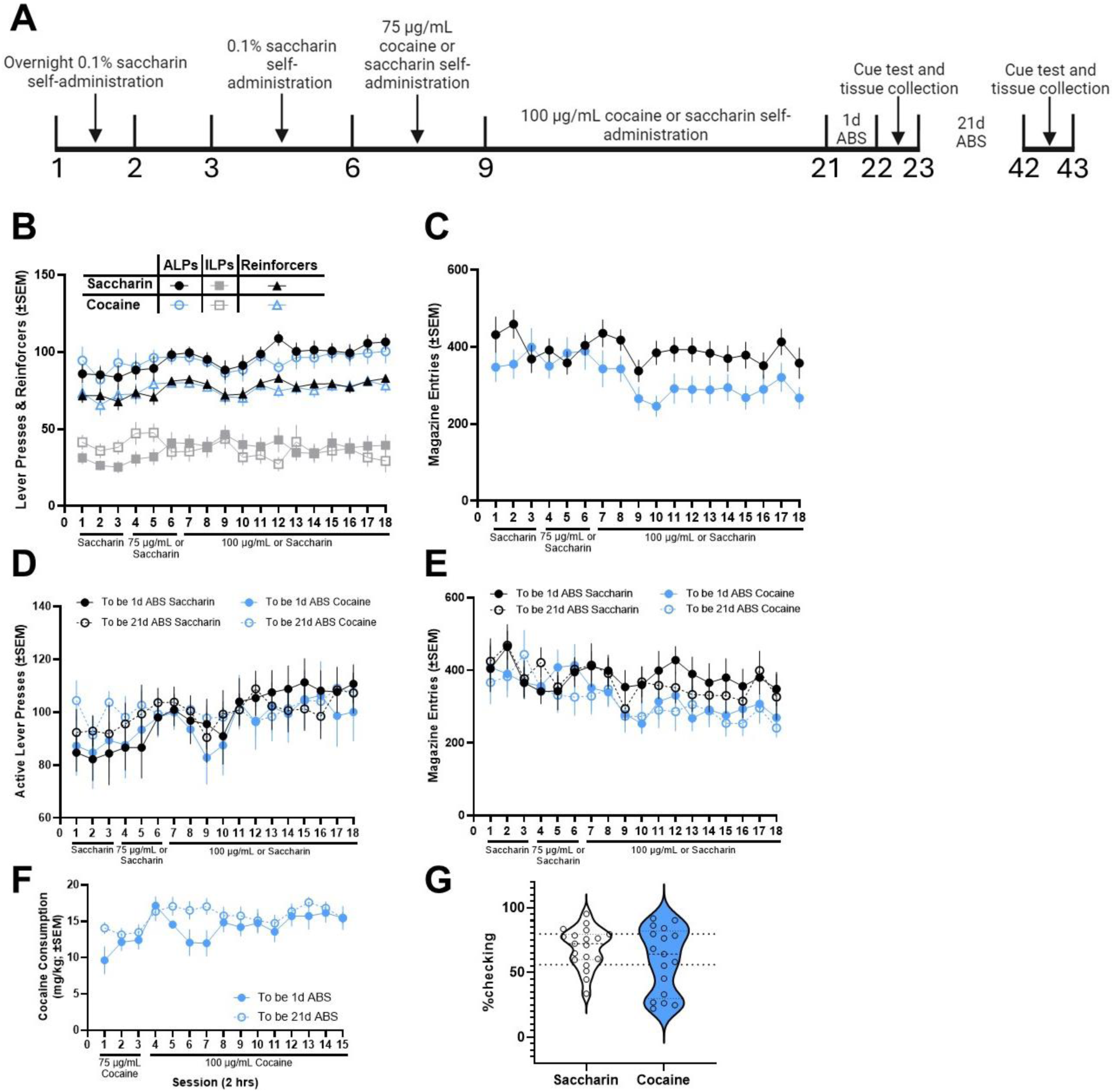
Cocaine and saccharin self-administration behavior prior to forced abstinence. **(A)** Mice were trained in an overnight session of 0.1% saccharin self-administration at 20 µL/reinforcer prior to self-administration of 50 µL/reinforcer of saccharin for 3 days. Mice then continued to self-administer saccharin or began self-administration of 75 µg/mL cocaine for 3 days followed by 100 µg/mL cocaine for 12 days. Mice then entered 1 or 21 days of forced abstinence prior to a cue test, where they were sacrificed immediately after testing for brain tissue collection. **(B-C)** Overall active and inactive lever presses, reinforcers earned, and magazine entries did not significantly differ between cocaine and saccharin self-administering animals. **(D-E)** When analyzed by future abstinence group assignment, active lever presses and magazine entries did not differ significantly as a function of abstinence for both cocaine and saccharin self-administering mice. **(F)** Among cocaine self-administering mice, cocaine consumption did not differ significantly between future abstinence groups. **(G)** Magazine checking behavior, defined as the percentage of active lever presses that were immediately followed by a magazine entry, did not significantly differ between cocaine and saccharin self-administering mice.

## Results

### Blood and brain cocaine concentrations following oral self-administration

We first sought to validate the concentrations of cocaine, and its major metabolite benzoylecgonine (BZE), within blood and brain tissue at two different time points (i.e., after 1 or 2 hrs of self-administration), using a within-subjects design for blood analyses (**Figure 1A**). Self-administration behavior for these animals is depicted in **Supplemental Figure S2**. Summary statistics of these analyte concentrations are summarized in **Tables 1 and 2**. Sufficient blood could not be collected for one animal at the 2 hr timepoint, so this mouse was excluded from the blood cocaine and BZE analyses here. A paired t-test revealed that blood concentrations of cocaine did not differ between 1 and 2 hrs of cocaine self-administration (*t*_(10)_ = 0.3589, p = 0.7271; **Fig 1B**). In contrast, a paired t-test revealed that blood BZE concentrations were significantly elevated after 2 hrs of cocaine self-administration compared to 1 hr of self-administration (*t*_(10)_ = 5.111, *p = 0.0005; **Fig 1C**). Simple linear regression analyses revealed that cocaine consumption significantly correlated with both blood cocaine (r^2^ = 0.2964, *p = 0.0072; **Fig 1D**) and BZE concentrations (r^2^ = 0.5823, *p < 0.0001; **Fig 1E**). Unlike blood cocaine concentrations, brain cocaine levels were significantly elevated after 2 hrs of cocaine self-administration compared to 1 hr of self-administration, as revealed by an unpaired t-test (*t*_(6)_ = 3.859, *p < 0.0084; **Fig 1F**). Brain BZE concentrations were also significantly elevated after 2 hrs of cocaine self-administration compared to 1 hr of self-administration as revealed by an unpaired t-test (*t*_(6)_ = 3.786, *p < 0.0091; **Fig 1G**). Simple linear regression analyses revealed a trend towards a significant correlation between cocaine consumption and brain cocaine levels (r^2^ = 0.4752, p = 0.0586; **Fig 1H**) and a significant correlation between cocaine consumption and brain BZE concentrations (r^2^ = 0.9268, *p = 0.0001; **Fig 1I**). Overall, these results indicate that blood cocaine levels within a 2 hr self-administration session achieve stability by at least halfway through the session, that cocaine is detectable in the brain, and that oral cocaine consumption is predictive of the levels of both cocaine and BZE within the blood and brain.

**Table 1.** Blood cocaine and BZE concentrations following 1 or 2 hours of oral cocaine self-administration.

| **Group** | **Mean Cocaine (ng/mL)** | **Cocaine S.D.** | **Mean BZE (ng/mL)** | **BZE S.D.** |
| --- | --- | --- | --- | --- |
| **1 hr S.A.** | 5.967 | 4.971 | 90.62 | 26.98 |
| **2 hr S.A.** | 7.145 | 5.933 | 797.6 | 449.7 |

**Table 2.** Brain cocaine and BZE concentrations following 1 or 2 hours of oral cocaine self-administration.

| **Group** | **Mean Cocaine (ng/mL)** | **Cocaine S.D.** | **Mean BZE (ng/mL)** | **BZE S.D.** |
| --- | --- | --- | --- | --- |
| **1 hr S.A.** | 1.6 | 1.364 | 6.9 | 2.634 |
| **2 hr S.A.** | 6.95 | 2.415 | 15.75 | 3.862 |

### Cocaine self-administration and cue-induced reward seeking

Here, mice underwent oral cocaine or saccharin self-administration prior to 1 or 21 days of forced abstinence and cue testing. A timeline of the behavioral paradigm is provided in **Figure 2A**. A repeated measures three-way ANOVA of active and inactive lever presses between cocaine and saccharin self-administering mice revealed a significant main effects of session (*F*_(17, 578)_ = 2.104; p = 0.0059) and lever (*F*_(0.4040, 13.74)_ = 241.5; p < 0.0001) as well as significant session X lever (*F*_(5.107, 173.6)_ = 2.984; p = 0.0125) and session X Reinforcer (*F*_(17, 578)_ = 2.769; p = 0.0002) interactions (**Fig 2B**). A repeated measures two-way ANOVA of magazine entries revealed a significant main effect of session (*F*_(17,578)_ = 3.478; p < 0.0001; **Fig 2C**) and trends towards a significant main effect of reinforcer (*F*_(1, 34)_ = 3.979; p = 0.0542) and session X reinforcer interaction (*F*_(17,578)_ = 1.639; p = 0.0503). Importantly, no significant main effect of abstinence group assignment was detected in active lever presses for saccharin (*F*_(1,17)_ = 0.2899; p = 0.5973) and cocaine (*F*_(1,15)_ = 1.816; p = 0.1978) (**Fig 2D**), magazine entries for saccharin (*F*_(1,17)_ = 0.1917; p = 0.6670) and cocaine (*F*_(1,15)_ = 1.017; p = 0.3291) (**Fig 2E**), and cocaine consumption among cocaine animals (*F*_(1,15)_ = 1.434; p = 0.2497) (**Fig 2F**) among mice that were later assigned to 1 or 21 days of forced abstinence (i.e., 1d ABS or 21d ABS). In addition to the overall macrostructure of self-administration behavior, we examined differences in magazine checking behavior (defined as the percentage of lever presses that were immediately followed by a magazine entry), which is a microstructural behavioral pattern that reflects tracking of response-outcome contingencies (Bryant et al., 2023). No significant difference in checking behavior was observed between saccharin and cocaine self-administering mice (*t*_(34)_ = 1.386; p = 1746; **Fig 3G**).

**Figure 3.**
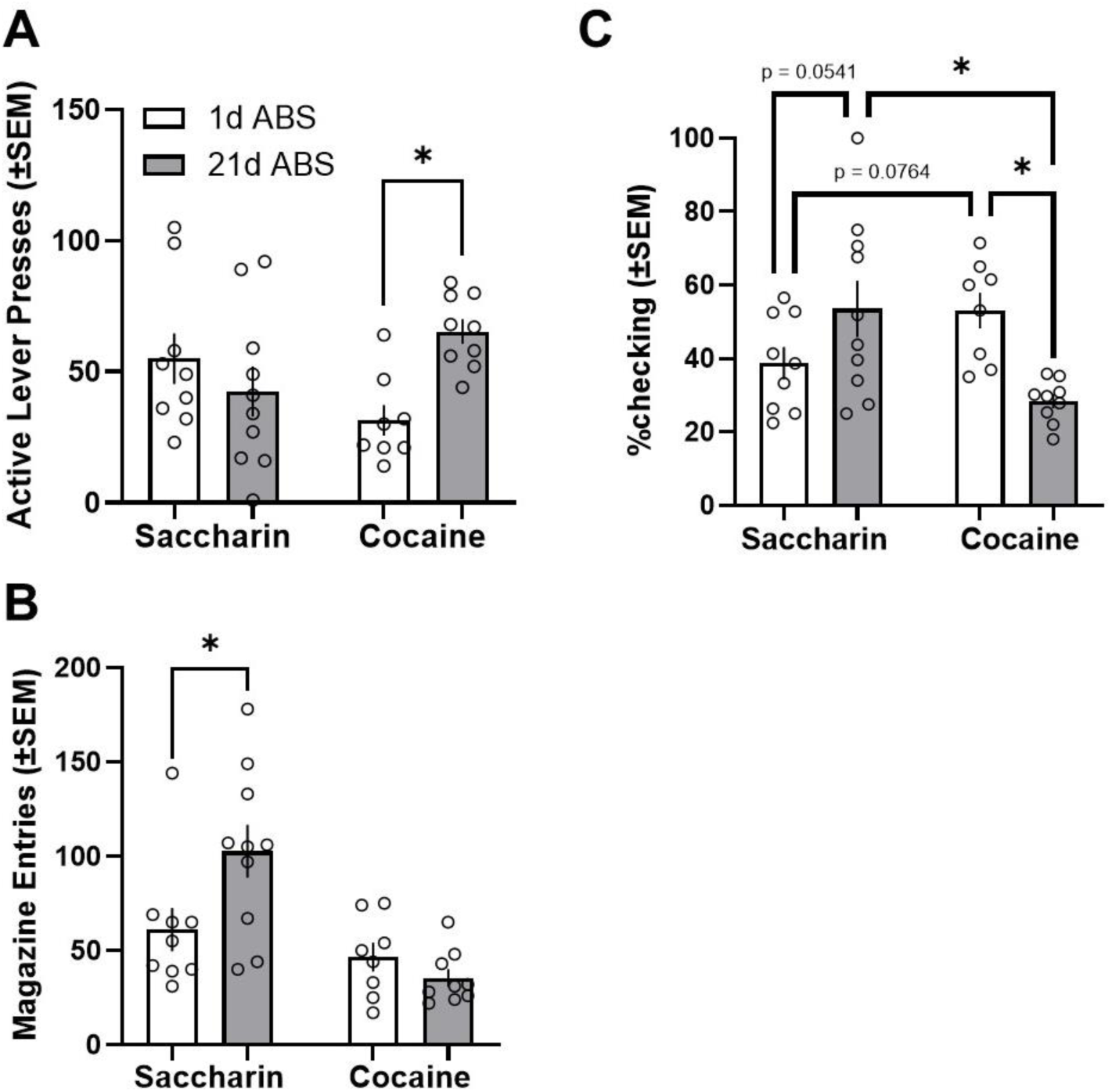
Incubation of cocaine and saccharin seeking and taking-related behavior following forced abstinence. Following forced abstinence, mice received a 1-hr cue test where active lever presses resulted in reward-paired cues but no reward. **(A)** Reward-seeking behavior, measured as active lever presses, was significantly increased after 21d abstinence (21d ABS) compared to 1d abstinence (1d ABS) for mice that self-administered cocaine but not saccharin. **(B)** Conversely, reward-consumption behavior, measured as magazine entries, was significantly increased after 21d ABS compared to 1d ABS for mice that self-administered saccharin but not cocaine. **(C)** Examining magazine checking behavior during the cue test, we observed a significant abstinence X reinforcer interaction effect, where checking behavior was significantly reduced from 1d ABS to 21d ABS among mice that self-administered cocaine. A trend towards an increase in checking behavior across abstinence was observed among mice that self-administered saccharin as well as a trend towards increased checking behavior among mice that self-administered cocaine compared to saccharin at 1d ABS. After 21d ABS, a significant reduction in checking behavior was observed among mice that self-administered cocaine compared to saccharin. *p < 0.05.

Following self-administration and forced abstinence, mice were tested for their active lever press and magazine entry responses during a cue test, where active lever presses resulted in cue presentation but no reward. A two-way ANOVA of active lever presses revealed a significant abstinence x reinforcer interaction effect (*F*_(1,32)_ = 8.398; p = 0.0067) and no significant main effects of abstinence or reinforcer. *Post-hoc* analyses revealed that among mice that self-administered cocaine, but not saccharin, a significant increase in active lever presses was observed at 21d ABS relative to 1d ABS (*p = 0.0128, **Fig 3A**). A two-way ANOVA of magazine entries revealed a significant abstinence x reinforcer interaction effect (*F*_(1,32)_ = 6.293; p = 0.0174), a significant main effect of reinforcer (*F*_(1,32)_ = 15.24; p = 0.0005), and no significant main effect of abstinence. *Post-hoc* analyses revealed that among mice that self-administered saccharin, but not cocaine, a significant increase in magazine entries was observed at 21d ABS relative to 1d ABS (*p = 0.0139, **Fig 3B**). Furthermore, a two-way ANOVA of magazine checking behavior revealed a significant reinforcer x abstinence interaction (*F*_(1,32)_ = 13.50; p = 0.0009; **Fig 3C**), and *post-hoc* analyses revealed a significant reduction in checking behavior from 1d to 21d ABS among mice that self-administered cocaine (*p = 0.0034) as well as a significant reduction in checking behavior at 21d ABS among mice that self-administered cocaine compared to saccharin (*p = 0.0018). Non-significant trends towards enhanced checking behavior at 1d ABS among mice that self-administered cocaine compared to saccharin (p = 0.0764) and enhanced checking behavior from 1d to 21 ABS among mice that self-administered saccharin (p = 0.0541) were also observed. These results suggest that the incubation response following oral self-administration and protracted abstinence is dependent on the reinforcer and the operant behavioral response being measured and that the behavioral microstructure of reward seeking following protracted abstinence is also reinforcer specific.

### Nucleus accumbens (NAc) expression of GLT-1 and glutamate receptors

Following the post-abstinence cue test, brains were harvested and P2 membrane fractions of the NAc were prepared to examine the expression of key glutamatergic substrates that are classically implicated in relapse-like cocaine seeking. A two-way ANOVA of GLT-1 expression revealed a significant main effect of reinforcer (*F*_(1,30)_ = 4.717; p = 0.0379) and no significant main effect of abstinence (*F*_(1,30)_ = 1.138; p = 0.2945) or abstinence X reinforcer interaction effect (*F*_(1,30)_ = 0.9132; p = 0.3469) (**Fig 4A**), where cocaine significantly reduced GLT-1 expression relative to saccharin regardless of abstinence. While simple linear regression analyses revealed no statistically significant correlations between GLT-1 expression and active lever presses or magazine entries for both cocaine (**Figs 4B-C**) and saccharin animals (**Figs 4D-E**), we did observe a trend towards a negative correlation between GLT-1 expression and both behaviors among mice that self-administered cocaine (r^2^ = 0.2257, p = 0.0735 for active lever presses; r^2^ = 0.2241, p = 0.0747 for magazine entries). Unlike GLT-1, two-way ANOVAs of GluA1 (**Fig 5A**) and GluA2 expression (**Fig 6A**) revealed no significant main effects of abstinence or reinforcer, nor was a significant abstinence X reinforcer interaction observed. Similarly, simple linear regression analyses revealed no significant correlations between GluA1 expression and active lever pressing or magazine entries for both cocaine (**Figs 5B-C**) and saccharin (**Figs 5D-E**). Such correlations were similarly not observed for GluA2 expression (**Figs 6B-E**). These results implicate glial glutamate transport as a neurobiological mechanism that is impaired by oral self-administration of cocaine.

**Figure 4.**
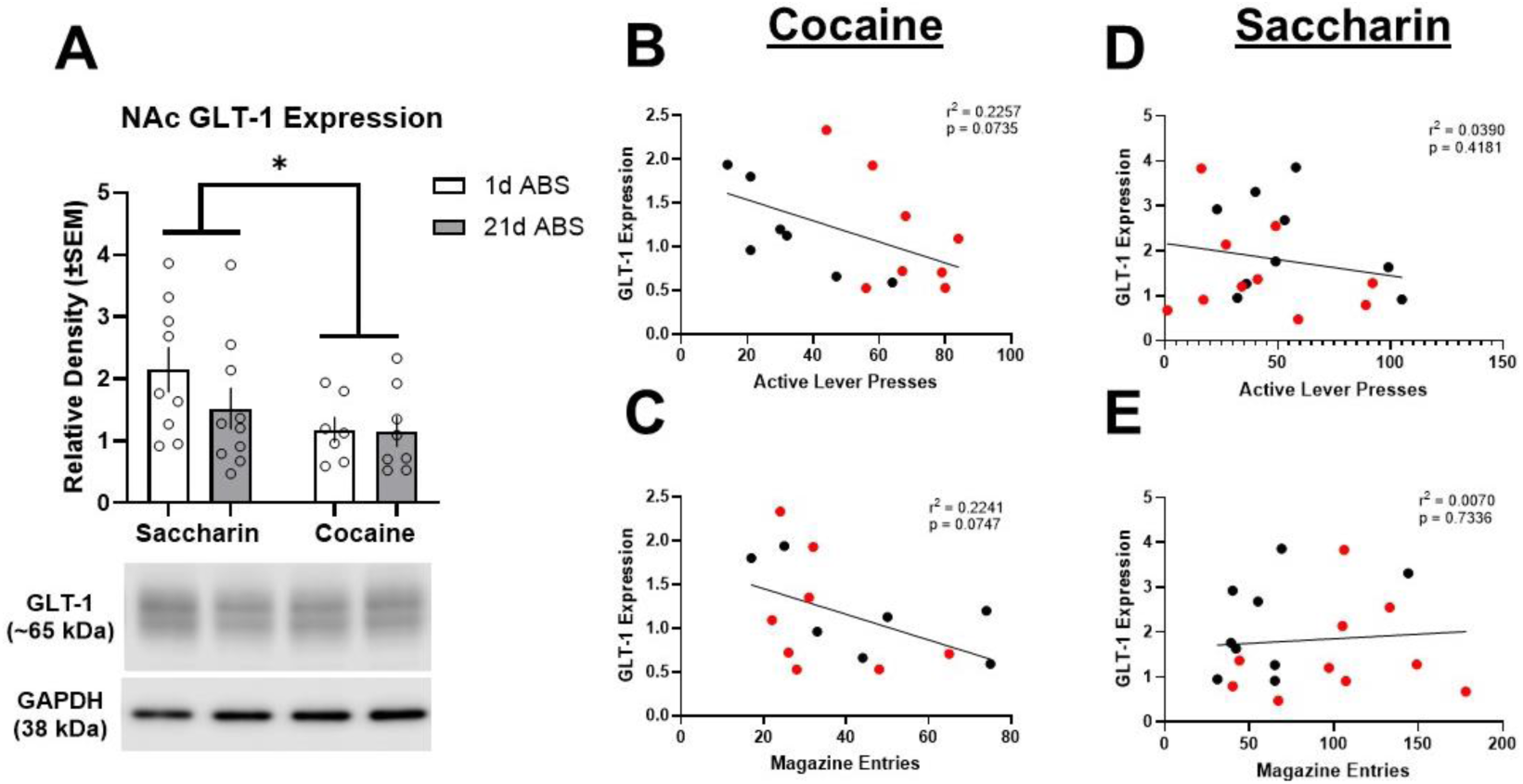
NAc GLT-1 expression following saccharin or cocaine self-administration and abstinence. P2 membrane fractions of the NAc were assessed for GLT-1 expression **(A)** A significant main effect of reinforcer was observed, where GLT-1 expression was suppressed within the NAc of mice that self-administered cocaine relative to saccharin. **(B-C)** A trend towards a negative correlation between active lever presses and GLT-1 expression (p = 0.0735), as well as between magazine entries and GLT-1 expression (p = 0.0747), was observed. **(D-E)** In contrast, no such trends towards a correlation between active lever presses or magazine entries and GLT-1 expression were observed. Black dots = 1d ABS. Red dots = 21d ABS. *p < 0.05.

**Figure 5.**
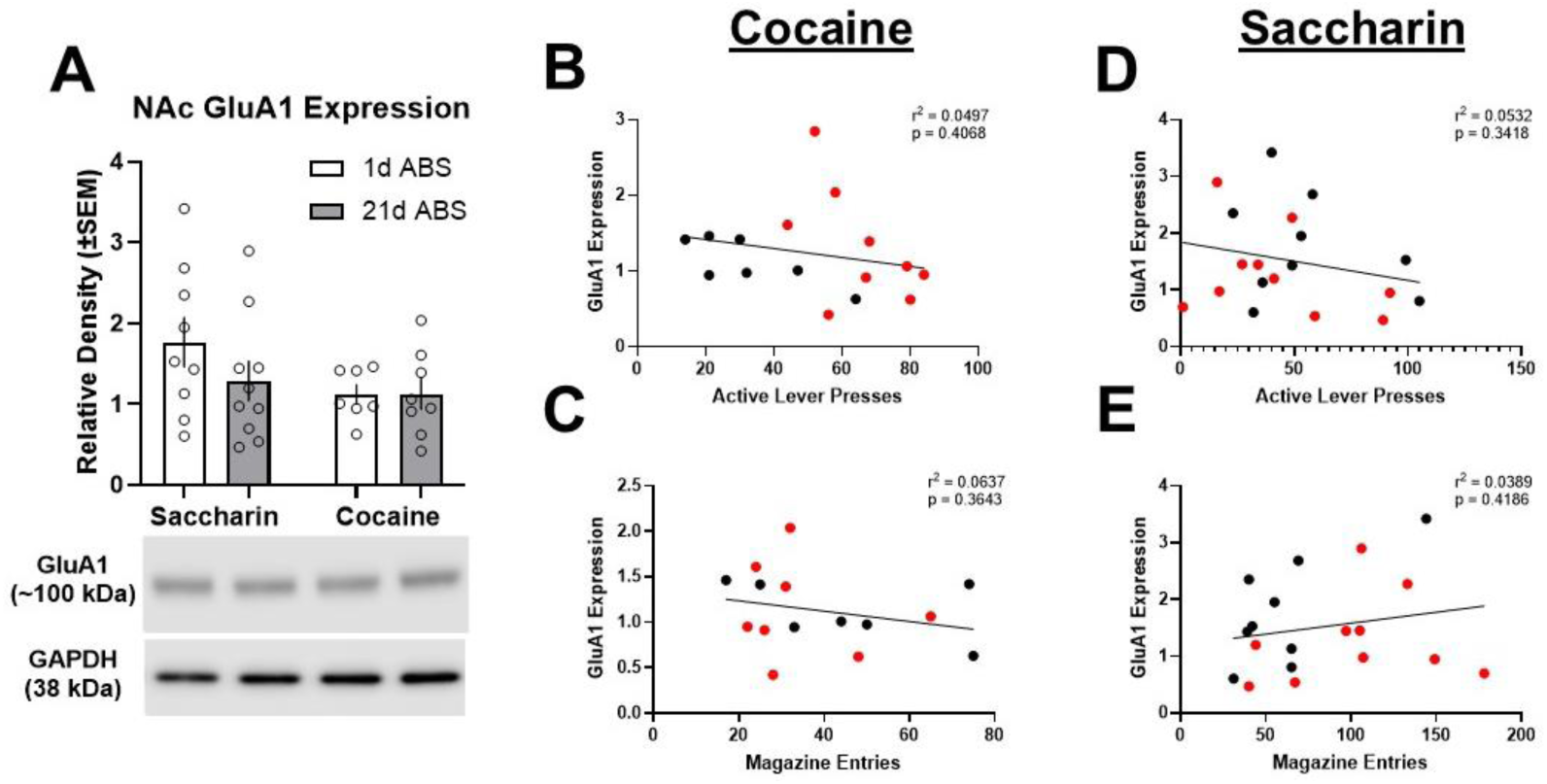
NAc GluA1 expression following saccharin or cocaine self-administration and abstinence. P2 membrane fractions of the NAc were assessed for GluA1 expression **(A)** No significant effects of abstinence or reinforcer on NAc GluA1 expression were observed. **(B-E)** No correlations were observed between active lever presses or magazine entries and GluA1 expression for either cocaine or saccharin. Black dots = 1d ABS. Red dots = 21d ABS. *p < 0.05.

**Figure 6.**
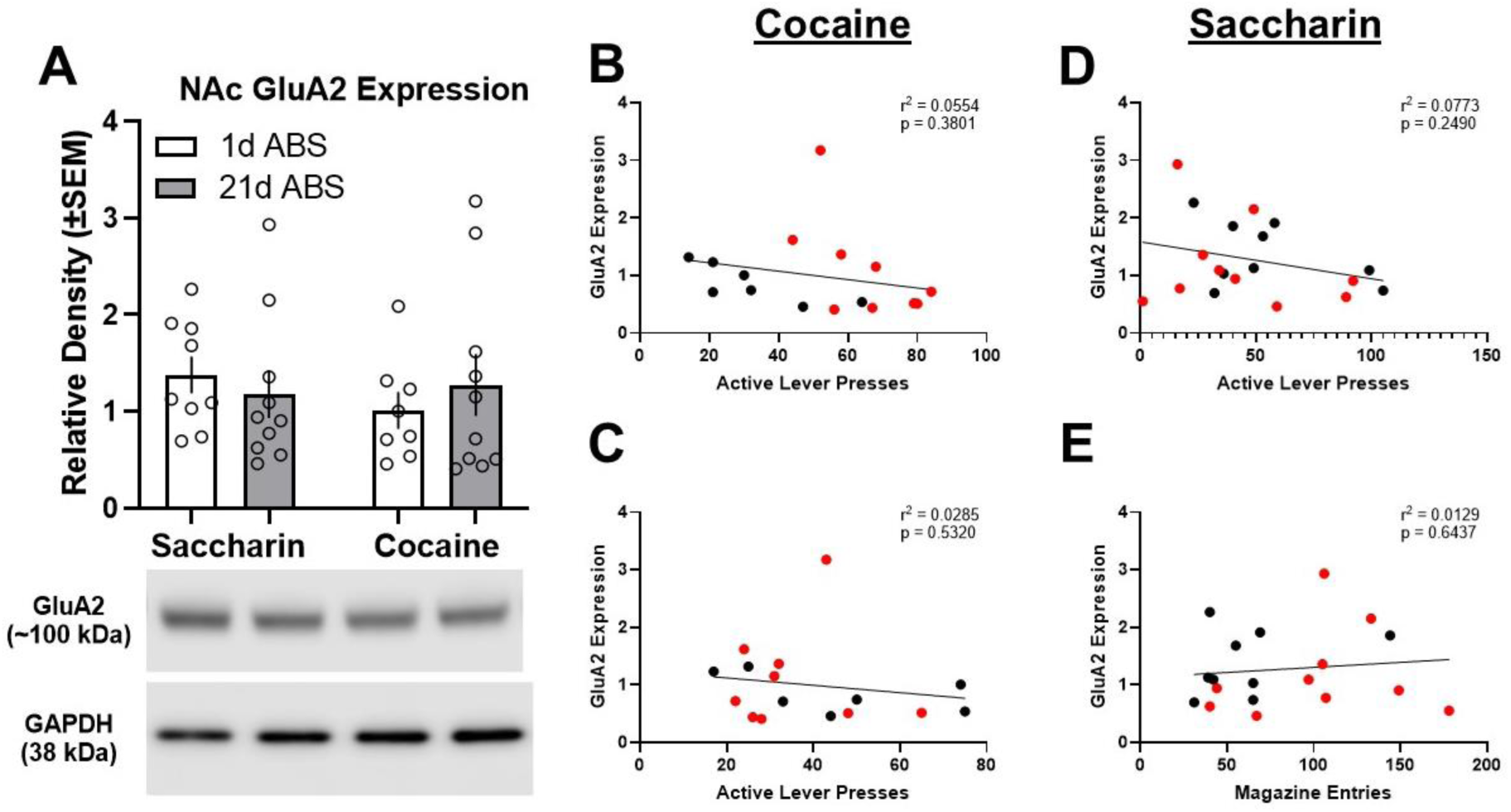
NAc GluA2 expression following saccharin or cocaine self-administration and abstinence. P2 membrane fractions of the NAc were assessed for GluA2 expression **(A)** No significant effects of abstinence or reinforcer on NAc GluA2 expression were observed. **(B-E)** No correlations were observed between active lever presses or magazine entries and GluA2 expression for either cocaine or saccharin. Black dots = 1d ABS. Red dots = 21d ABS.

## Discussion

There remains a dearth of studies that characterize the incubation of drug seeking using non-intravenous models of operant drug self-administration. Considering nearly all studies examining the incubation effect utilize forms of intravenous self-administration that do not dissociate appetitive from consummatory behavior, our current understanding of the neurobehavioral mechanisms of the incubation effect is incomplete. Thus, we sought to interrogate the neurobehavioral sequelae of oral cocaine self-administration and the incubation of cocaine seeking in this model. We demonstrate that mice will operantly respond for an oral solution of cocaine and express incubated cocaine seeking in a manner that is distinct from incubated saccharin seeking. Moreover, oral cocaine self-administration suppressed membrane expression of the glutamate transporter GLT-1 within the NAc, which is a key glutamatergic substrate implicated in cue-induced drug relapse. Taken together, these results highlight the utility of oral cocaine self-administration to study relapse-like behavior and its neurobiological correlates.

### The behavioral macro- and microstructures of oral cocaine self-administration and incubated reward seeking

The present study builds upon recent studies that have developed protocols for studying the neurobehavioral sequelae of oral cocaine self-administration in both adolescent and adult mice (Depoy et al., 2016; Li et al., 2022). Here, we confirm that mice will readily self-administer cocaine orally in a 0.1% saccharin solution, that cocaine and its major metabolite, BZE, are detectable in the brain and periphery following self-administration, and that oral cocaine consumption is predictive of both blood and brain levels of cocaine and BZE. Overall self-administration, as well as checking behavior, was similar between cocaine and saccharin. While not statistically significant, there was a trend towards reduced overall magazine entries towards the latter half of cocaine self-administration sessions compared to saccharin self-administration. A cocaine-induced change in checking behavior was observed during the post-abstinence cue test. Checking behavior observed during the cue test, defined as the percentage of active lever presses that were immediately followed by a magazine entry, can be described as a behavioral sequence that reflects continual monitoring and sampling of the response-outcome contingency between the active lever and magazine (Bryant et al., 2023). Previous work demonstrates that prolonged cocaine self-administration, whether oral or intravenous, can produce insensitivity to response-outcome devaluation in a manner that some argue reflects a shift from goal-directed to habitual behavior (Tiffany, 1990; Miles et al., 2003; Zapata et al., 2010), though that was not assessed in the current study. Whether or not the reduced checking behavior during the cue test reflects a shift towards habitual behavior, these data suggest that certain cocaine-induced adaptations in behavioral microstructures, such as magazine checking, may be dependent on abstinence and, perhaps, the route of administration and subsequent pharmacokinetic factors.

Increased cocaine craving and seeking across a protracted period of abstinence, known as the incubation of drug craving, is a key addiction-related process that has been validated across various animal models of SUDs. However, one limitation of these preclinical studies is that they predominantly employ intravenous routes of drug self-administration (Pickens et al., 2011; Li et al., 2015), where appetitive and consummatory behavior are typically conflated and inherently not dissociable because these two behaviors occur on the same operandum (Roberts et al., 2013). Thus, one strength of the model presented here is that lever pressing for reward is a distinct action that is separate from the retrieval and subsequent consumption of said reward. Here, we show that mice with a history of cocaine self-administration will exhibit an incubation of cocaine seeking at the active lever that delivers cocaine-paired cues. This is in contrast to mice with a history of saccharin self-administration, where incubation was observed at the magazine, where mice would otherwise retrieve and consume reward. These results parallel findings from a recent study that trained rats to self-administer cocaine or saccharin in a “seeking-taking” paradigm, where lever pressing on a FI schedule (i.e., “seeking” lever) resulted in presentation of a lever that delivered reward on a FR schedule (i.e., “taking” lever) (Leonard et al., 2024). Here, incubation of FI lever responses were observed for cocaine but not saccharin. These past and current data suggest that the route of administration, the reinforcer type, and perhaps the schedule of reinforcement are likely important variables that collectively influence the macro-and microstructure of incubated reward seeking following protracted abstinence.

### NAc GLT-1, but not AMPA receptor subunit, expression is impaired by oral self-administration of cocaine

A key finding in this study that mirrors the results of intravenous cocaine self-administration studies is the cocaine-induced downregulation of NAc GLT-1 expression. Intravenous self-administration of cocaine impairs GLT-1 function and expression within the NAc, and suppression of cue-induced reinstatement of cocaine seeking by drugs such as the beta lactam antibiotic ceftriaxone and the antioxidant *N*-acetylcysteine requires restoration of GLT-1 expression (Knackstedt et al., 2010; Trantham-Davidson et al., 2012; Reissner et al., 2015; LaCrosse et al., 2016). Importantly, this phenomenon extends to other addictive drugs. For example, we have shown that inhibition of cue-induced nicotine self-administration by *N*-acetylcysteine requires the rescue of GLT-1 expression (Namba et al., 2020), and other studies have demonstrated that drugs such as alcohol and opioids similarly impair striatal GLT-1 expression (Alhaddad et al., 2014; Shen et al., 2014). While not statistically significant, we observed a trend towards a negative correlation between GLT-1 expression and both active lever presses and magazine entries during the cue test. This trend was specific to cocaine, with no such trends observed for saccharin seeking. Indeed, this corroborates the plethora of preclinical studies, across addictive drugs, that implicate impaired GLT-1 expression within the NAc as a key drug-induced glutamatergic adaptation that contributes to relapse vulnerability. Nevertheless, it is possible we would have seen a more pronounced effect on GLT-1 expression and its correlation with cocaine seeking with a longer withdrawal period and/or extended access cocaine self-administration as some previous studies suggest (Fischer-Smith et al., 2012; Fischer et al., 2013; Kim et al., 2018).

While enhanced NAc expression of the GluA1 AMPA receptor subunit was an expected neural correlate of incubated cocaine seeking, we did not observe this. Increased NAc expression of GluA1 is a glutamatergic neuroadaptation that can facilitate postsynaptic excitability of medium spiny neurons within the NAc and cocaine relapse-like behavior, and pharmacological inhibition of GluA2-lacking calcium-permeable AMPARs (CP-AMPARs) within the NAc suppresses the incubation of cocaine seeking (Conrad et al., 2008; Purgianto et al., 2013). Similar to GLT-1, some studies suggest that cocaine-induced adaptations to AMPA receptor subunit composition (particularly an increase in GluA2-lacking, calcium-permeable AMPA receptors) are much more pronounced following extended access to cocaine (e.g., 6 hr access) and/or later withdrawal timepoints (e.g., 40+ days) (Conrad et al., 2008; Fischer-Smith et al., 2012; Fischer et al., 2013; Kim et al., 2018; Christian et al., 2021). Nevertheless, other studies have confirmed cocaine-induced changes in GLT-1 and surface expression of GluA1 within the NAc following short access to cocaine self-administration and two weeks of withdrawal, albeit under extinction conditions (Wolf and Tseng, 2012; Reissner et al., 2015; LaCrosse et al., 2016).

## Limitations & Future Directions

There are some limitations and potential future directions to this study that are worth noting. Firstly, this study only used male mice. This initial study sought to first characterize and validate the oral self-administration model prior to interrogating its neurobehavioral effects on incubated reward seeking. However, sex differences in cocaine self-administration and the incubation of cocaine seeking are well-documented (Hu et al., 2003; Swalve et al., 2015; Corbett et al., 2021; Towers et al., 2023). While very little is known regarding sex differences in the oral self-administration of cocaine, a recent report demonstrated that sex is an important biological determinant of future habit behaviors among animals that escalate oral cocaine seeking (Depoy et al., 2016), and women are more likely than men to develop cocaine dependence even though more men use cocaine (O’Brien and Anthony, 2005). Thus, examination of female animals is a crucial next step to advance this model.

Another limitation of the present study is that we only examined the impact of short access self-administration and 21 days of forced abstinence on incubated reward seeking and associated neuroadaptations within the NAc. As discussed above, certain cocaine-induced glutamatergic adaptations are more pronounced with extended access self-administration and/or longer abstinence periods (e.g., 40+ days). Indeed, future studies implementing the model presented here may consider examining both extended access self-administration conditions and/or extended withdrawal timepoints, as they may yield more pronounced effects on glutamatergic substrates as well as more divergent behavioral patterns of responding during self-administration and cued seeking following abstinence. However, it is important to note that traditional extended access conditions do not necessarily recapitulate the temporal patterns of cocaine use observed in humans, which also more often than not occur in the context of polysubstance use (PSU; Liu et al., 2018; Fitzgerald et al., 2024). We argue that future studies that holistically model factors such as route of administration, PSU, order of drug administration in the case of PSU, and frequency of drug use (e.g., continuous versus intermittent access) are more likely to successfully model the neurobehavioral sequelae of cocaine use disorder in humans.

Lastly, it is worth considering that this model is limited to the administration of low unit doses of cocaine. Due to the inherently bitter and aversive taste of cocaine, as well as the adverse gastrointestinal and hepatic effects of acute, high doses of orally administered cocaine (Farrar and Kearns, 1989; Mehanny and Abdel-Rahman, 1991; Labib et al., 2001), we were limited in how concentrated a unit dose of cocaine could be when combined with 0.1% saccharin. Other studies utilizing alternative vehicles, such as a sucrose solution (Depoy et al., 2016; Li et al., 2022), could potentially help ameliorate this constraint on cocaine dosing. While the need for a vehicle such as sucrose or saccharin to overcome the aversive taste of cocaine is an inherent limitation here, we contend that the differential effects on incubated seeking behavior and the parallel neurobiological effects on GLT-1 expression between saccharin and cocaine in this model highlight its utility and concurrent validity when compared to more traditional self-administration models. Moreover, as referenced above, cocaine is rarely consumed in the absence of other reinforcers (e.g., alcohol). Thus, understanding the neurobehavioral effects of cocaine in the context of concurrent use of other reinforcers is highly warranted.

## Conclusions

In conclusion, we demonstrate that adult male mice that orally self-administer cocaine exhibit a pattern of incubated seeking behavior that is distinct from that of saccharin self-administering mice. Moreover, we demonstrate that GLT-1, which is a key glutamatergic substrate that is classically implicated in cocaine relapse, is downregulated by oral self-administration of cocaine. These results highlight the utility of oral, operant self-administration of cocaine as a viable alternative to intravenous self-administration when studying the incubation of craving effect in preclinical rodent models, particularly when practical constraints limit the use of intravenous catheters for self-administration. Future studies examining these effects in females, utilizing alternative drug access schedules and withdrawal timepoints, and building in polysubstance use into the model are all warranted.

## Acknowledgements

We would like to thank staff members of the Center for Forensic Science Research & Education and members of the Barker Lab for their collaboration and excellent technical expertise on this project.

## Author Contributions

*Mark D Namba* – Conceptualization, Formal Analysis, Funding Acquisition, Methodology, Investigation, Validation, Visualization, Writing - Original Draft Preparation, and Writing - Reviewing & Editing; *Samuel L Goldberg; Christine M Side; Christina M Curran-Alfaro; Abhiram Yadlapalli* – Investigation and Writing - Reviewing & Editing. *Melissa F Fogarty; Amanda LA Mohr; Devin Kress* – Investigation, Validation, Methodology, Resources, and Writing - Reviewing & Editing. *Jacqueline M Barker* – Conceptualization, Formal Analysis, Funding Acquisition, Methodology, Investigation, Validation, Visualization, Project Administration, Resources, Supervision, Writing - Original Draft Preparation, and Writing - Reviewing & Editing.

## Data Availability

All data reported in the manuscript are secured on Drexel University cloud servers under controlled access and will be made available upon request. Requests should be made to Dr. Jacqueline Barker.

---

**Supplemental Table 1.** Method Gradient.

| **Supplemental Table 1.** |  |
| --- | --- |
| **Method Gradient** |  |
| **Time** | **%MPB** |
| 0 | 10 |
| 4 | 90 |
| 4.10 | 10 |
| 5 | 10 |

**Supplemental Table 2.** Antibody concentrations for western blots.

| **Supplemental Table 2. Antibody concentrations for western blots** |  |  |
| --- | --- | --- |
| **Target** | **1°Ab Concentrations** | **2° Ab Concentrations** |
| **GluA1** | 1:1,000; Abcam ab109450 | 1:5,000 anti-Rb HRP; Abcam ab97080 |
| **GluA2** | 1:1,000; Abcam ab206293 | 1:5,000 anti-Rb HRP; Abcam ab97080 |
| **GLT-1** | 1:3,000; Abcam ab41621 | 1:30,000 anti-Rb HRP; Abcam ab97080 |
| **GAPDH** | 1:3,000; Cell Signaling D16H11 | 1:30,000 anti-Rb HRP; Abcam ab97080 |

**Supplemental Figure S1.**
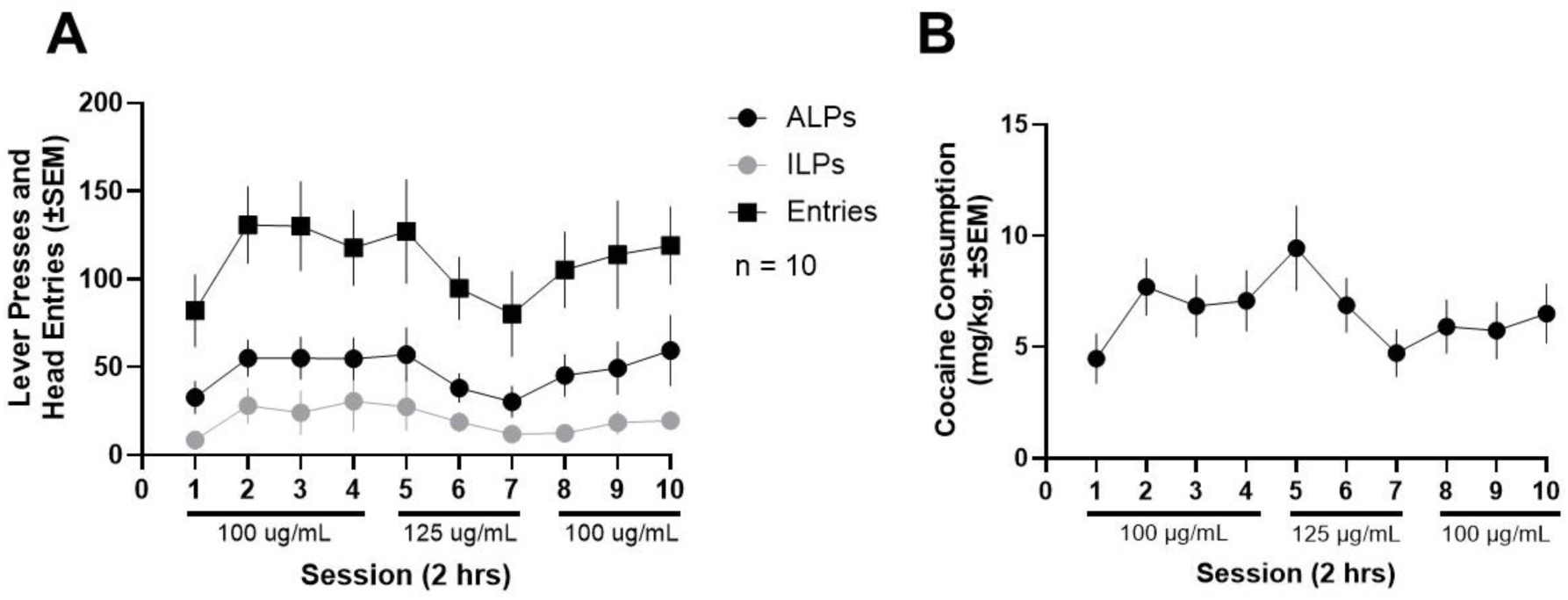
Mice achieve stable oral self-administration of cocaine at 100 µg/mL. We observed with a pilot cohort of mice (n=10) that stable cocaine consumption could be achieved at a concentration of 100 µg/mL in a 0.1% saccharin solution. However, increasing this concentration to 125 µg/mL did not result in an increase in (**A**) cocaine self-administration behavior and (**B**) cocaine consumption.

**Supplemental Figure S2.**
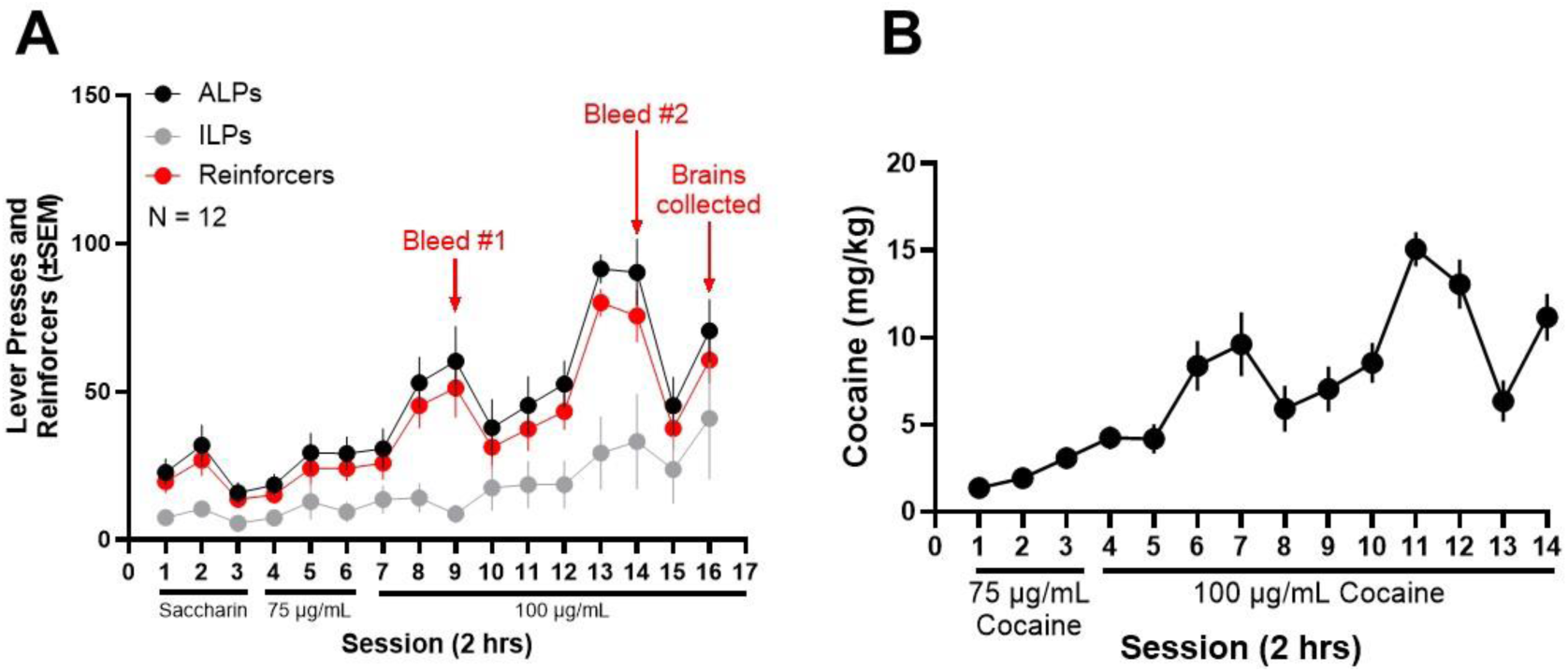
Cocaine self-administration behavior for blood and brain cocaine and BZE experiment. Using a within-subjects design, submandibular blood was collected after 1 or 2 hours of self-administration for all mice (n=12) at sessions 3 and 8 of 100 µg/mL cocaine self-administration. Mice were sacrificed for brain tissue collection at session 10. **(A)** Active and inactive lever presses and reinforcers earned across self-administration. **(B)** Cocaine consumption across self-administration.

